# BaseNet: A Transformer-Based Toolkit for Nanopore Sequencing Signal Decoding

**DOI:** 10.1101/2024.06.02.597014

**Authors:** Qingwen Li, Chen Sun, Daqian Wang, Jizhong Lou

**Affiliations:** Key Laboratory of Epigenetic Regulation and Intervention, Center for Excellence in Biomacromolecules, Institute of Biophysics, Chinese Academy of Sciences, Beijing 100101, China; University of Chinese Academy of Sciences, Beijing 100049, China; Beijing Polyseq Biotech Co. Ltd., Beijing 100089, China

**Keywords:** nanopore sequencing, basecall, transformer, machine learning algorithm

## Abstract

Nanopore sequencing provides a rapid, convenient and high-throughput solution for nucleic acid sequencing. Accurate basecalling in nanopore sequencing is crucial for downstream analysis. Traditional approaches such as Hidden Markov Models (HMM), Recurrent Neural Networks (RNN), and Convolutional Neural Networks (CNN) have improved basecalling accuracy but there is a continuous need for higher accuracy and reliability. In this study, we introduce BaseNet, an open-source toolkit that utilizes transformer models for advanced signal decoding in nanopore sequencing. BaseNet incorporates both autoregressive and non-autoregressive transformer-based decoding mechanisms, offering state-of-the-art algorithms freely accessible for future improvement. Our research indicates that cross-attention weights effectively map the relationship between current signals and base sequences, joint loss training through adding a pair of forward and reverse decoder facilitate model converge, and large-scale pre-trained models achieve superior decoding accuracy. This study helps to advance the field of nanopore sequencing signal decoding, contributes to technological advancements, and provides novel concepts and tools for researchers and practitioners.

## Introduction

Nanopore sequencing has emerged as a pivotal technology in modern genomics, offering fast, convenient and high-throughput sequencing for both DNA and RNA. This technique involves passing a DNA or RNA strand through a nanopore and measuring changes in ionic current to determine the sequence of nucleotides. Due to its speed, cost-effectiveness, and portability, nanopore sequencing has been widely applied in genomics research, medical diagnostics, personalized medicine, and other related fields ^[1–7]^.

Accurate decoding of nucleic acid sequences from the current signal change is crucial for effective downstream analysis in nanopore sequencing. These signal changes are complex, necessitating the use of artificial intelligence-based algorithms to decode the base sequence accurately ^[8]^. However, raw electrical signals are often influenced by various factors including system noise, base drift, and the heterogeneity of through-pore velocities etc. Consequently, achieving high-precision signal decoding in nanopore sequencing is a challenging yet essential for its widespread application and reliability ^[9]^.

Previous studies have predominantly employed basecallers constructed using methods such as Hidden Markov Models (HMM) ^[10]^, Recurrent Neural Networks (RNN), and Convolutional Neural Networks (CNN) ^[11–13]^ to decode the raw electrical signals in nanopore sequencing. While these methods have achieved significant success, they also have limitations. HMMs require manual feature extraction from signals and have limited capabilities for modeling long-range dependencies ^[14]^. RNNs can model long-range dependencies but suffer from issues such as vanishing and exploding gradients. CNNs, on the other hand, struggle to capture temporal information in sequences. These limitations restrict the decoding accuracy and reliability of some existing basecallers.

To address these challenges, the transformer model was originally proposed for machine translation tasks ^[15]^, leveraging self-attention and multi-head attention mechanisms to capture long-range dependencies in sequences. It has since achieved remarkable milestones in natural language processing, image recognition ^[16–18]^, and automatic speech recognition ^[19, 20]^, demonstrating powerful capabilities in biology as well, including protein generation, structure prediction, drug design, and biological sequence recognition ^[21–23]^. Transformer’s ability to effectively model long-range dependencies makes it a promising tool for many applications. However, in the field of nanopore sequencing, the application and development of transformer remains limited. Recent efforts have only utilized the transformer encoder and connectionist temporal classification (CTC) decoder for basecalling ^[24]^, leaving much potential for further exploration and advancement.

Here, we introduce BaseNet, an open-source toolkit that provides a transformer-based advanced algorithm platform for signal decoding in nanopore sequencing. BaseNet features several key components: 1). Autoregressive decoding: a transformer model using beam search for enhanced accuracy; 2). Non-autoregressive decoding: a transformer with a rescore decoding mechanism, trained using a combination of CTC and attention-based encoder-decoder (AED) ^[25]^; 3). Paraformer: a non-autoregressive decoder employing a Continuous Integrate-and-Fire (CIF) based predictor and a glancing language model (GLM) based generator ^[26]^; 4). Large-scale pre-trained model: a model fine-tuned using contrastive learning and diversity learning for improved performance on nanopore sequencing data ^[27]^; 5). Conditional random field (CRF) model: refined by a linear complexity attention mechanism to enhance decoding efficiency ^[28]^.

To our knowledge, BaseNet is the first to introduce transformer-based autoregressive and non-autoregressive decoding mechanisms in nanopore sequencing. Importantly, BaseNet provides several state-of-the-art transformer algorithms publicly available for for free access and further improvement. Below we outlines the main algorithm principles and model structures within BaseNet, as well as experimental results on key benchmark datasets of nanopore sequencing. Our results demonstrates that BaseNet achieves competitive performance, comparable to recent basecallers from Oxford Nanopore Technologies (ONT). The cross-attention weights in the transformer model effectively map the relationship between current signals and base sequences, joint loss training by adding forward and reverse decoders on the top of model improves convergence, and large-scale pre-trained model achieves higher decoding accuracy. Our findings also reveal the existence of common ‘generic features’ between speech waveforms and sequencing signals in model representation, highlighting the versatility and robustness of BaseNet in handling diverse signal types.

Thus, our studies advance the development of nanopore sequencing signal decoding, contribute to technological improvement, and provide novel ideas and valuable references for the field.

## Materials and methods

The raw current signals generated by nanopore sequencing are fed into BaseNet, which decodes the base sequences using its advanced decoding methods.

### Benchmark dataset

The benchmark dataset used in this study was proposed by Wick et al. and has been widely adopted for training and testing multiple basecallers^[29]^. It consists of both a training set and a test set. The training set includes genomes from fifty species, comprising 30 Klebsiella pneumoniae genomes, 10 Enterobacteriaceae genomes, and 10 Proteobacteria genomes. The test set contains genomes from 10 species, including 4 Klebsiella pneumoniae genomes, as well as genomes from Acinetobacter pittii, Haemophilus haemolyticus, Serratia marcescens, Shigella sonnei, Staphylococcus aureus, and Stenotrophomonas maltophilia. Each dataset includes raw current signals along with their corresponding contig sequences. To ensure the accuracy of the label sequences, Wick et al. performed second-generation sequencing and assembly to obtain the contig sequences.

Due to the long read lengths in nanopore sequencing, each current signal is exceptionally lengthy. For the model’s training and validation, we divided the original current signals into matrices with a maximum length of 10,000 sampling points to enhance training efficiency. To ensure accurate base sequences for each fragment and effective training, we first performed basecalling on the electrical signals using Guppy^[30]^. The resulting sequences were then aligned to the reference genome, and only when mapping the alignment accuracy exceeded 60%, the signal and its corresponding reference genome segment were retained for the training and validation sets.

### Self-attention and fast-attention mechanism

The self-attention mechanism enables the model to simultaneously attend to all positions in the sequence, capturing global dependencies. This mechanism also automatically calculates importance weights for each position based on the input sequence, allowing the model to focus on meaningful positions relevant to the current task. Unlike traditional fixed-weight mechanisms, self-attention provides greater flexibility to adapt to various tasks and inputs. The computation is as follows:

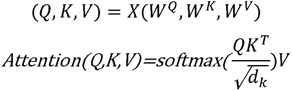

Here, *X* is the input matrix, (*W^Q^*, *W^K^*, *W^V^*) are three trainable parameter matrices. *Q*, *K*, and *V* are the query, key, and value matrices, respectively, obtained by linear transformations of the input matrix. *d*_k_ denotes the dimension of the key.

The multi-head self-attention mechanism allows the model to learn multiple different attention representations simultaneously. Each attention head can focus on different parts of the sequence, providing independent expressive capabilities. This approach captures semantic information at various levels, enhancing the model’s ability to comprehend the input sequence and improve its representation and generalization capabilities. And the computation follows:

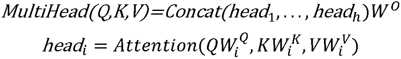

Where *W*^O^ is a trainable parameter matrix.

The algorithmic complexity of the attention mechanism described above is *O*(*N*^2^·*d*), where *N* represents the sequence length and *d* denotes the dimensionality of the matrices, resulting in a quadratic increase in computational efficiency relative to the sequence length. To alleviate computational costs, Wu et al. proposed a fast attention mechanism with linear complexity^[28]^. This method involves training separate matrices *α* and *β*, both with dimensions [*N*, 1], for *Q* and *K*, respectively. These matrices are used to perform a weighted summation on *Q* and *K*, resulting in transformed matrices *Q*’ and *K*’. Subsequently, the transformed matrices are multiplied together, reducing the complexity to *O*(*N*·*d*)^[28]^. The fast attention mechanism we have developed is illustrated in Figure 1.

**Figure 1.**
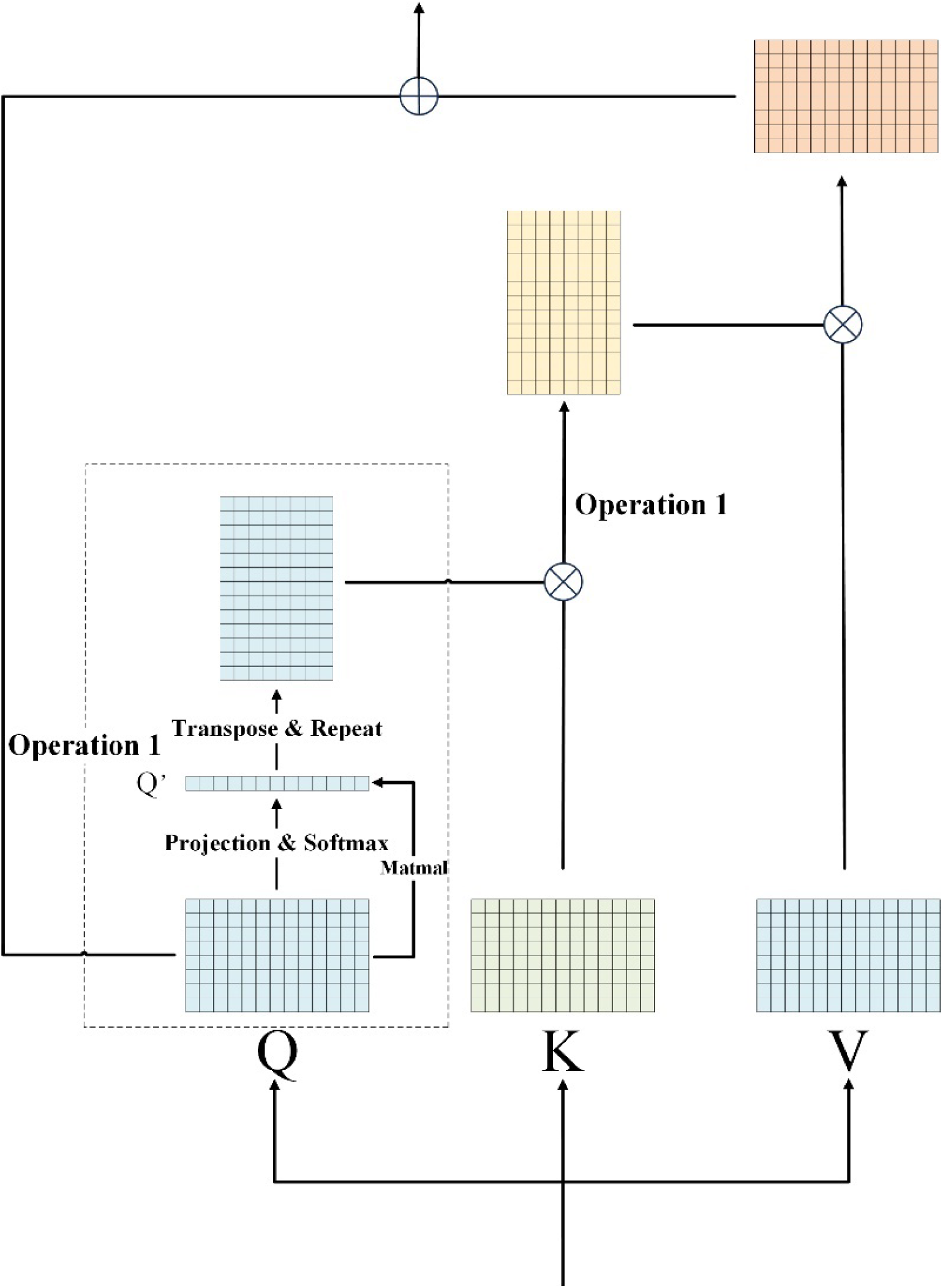
Fast attention mechanism in BaseNet. This schematic illustrates the fast attention mechanism used in BaseNet. The method involves training separate matrices, *α* and *β*, each with dimensions [*N*, 1], for *Q* and *K*, respectively. These matrices are used to perform a weighted summations on *Q* and *K*, resulting in transformed matrices *Q*’ and *K*’. The transformed matrices are then multiplied together, reducing the computational complexity to *O*(*N*·*d*).

### Autoregressive transformer and beam search

The model in this study draws inspiration from the transformer model initially proposed by the Google Machine Translation team^[15]^, with specific adaptations tailored for nanopore sequencing data as shown in Figure 2. It primarily comprises convolutional modules for feature extraction and down-sampling, an encoder for context modeling to generate hidden vectors, and a decoder for predicting the next time step output based on hidden vectors and generated sequences. Both the encoder and decoder modules consist of 8 layers. The model employs the self-attention mechanism in both the encoder and decoder stages.

**Figure 2.**
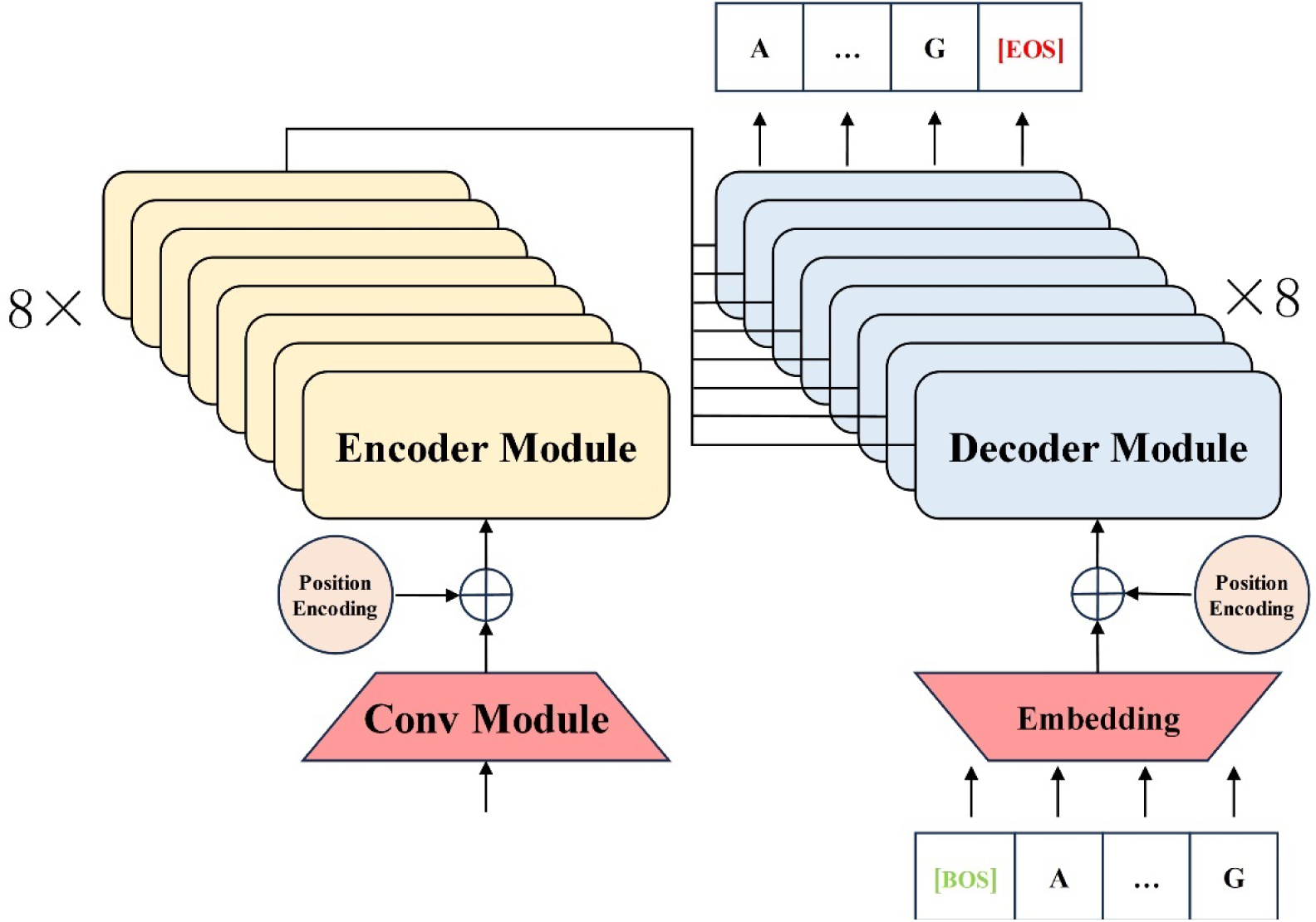
Autoregressive Transformer-based model architecture for nanopore sequencing. The schematic illustrates the architecture of the autoregressive transformer model tailored for nanopore sequencing data. The model includes convolutional modules for feature extraction and down-sampling, an encoder composed of 8 layers for context modeling, and a decoder with 8 layers for sequence generation.

In the inference stage, to enhance prediction accuracy, we combine autoregressive decoding with a beam search strategy. A beam size of 4 is selected, meaning that at each time step *t* during inference, the model retains the top 4 sequences with the highest scores. These sequences are then used as input for predicting the sequences at time step *t*+1 (i.e., retains top sequences with the highest scores among the 16 candidate sequences). The process is repeated iteratively until the decoding is complete. Finally, the sequence with the highest cumulative score across all time steps is selected as the decoding result.

### Pre-training and fine-tuning based on large-scale model

Baevski et al. previously introduced wav2vec2.0 ^[27]^, a self-supervised pre-training method that has shown significant success in speech recognition. Inspired by their research, we develops a large-scale model based on contrastive learning and diversity learning (Figure 3). The model includes a feature extraction module (comprising 7 1D convolution layers), an encoder module for context modeling (consisting of 12 transformer encoding layers), and a quantization module for learning discrete common features (implemented using a linear layer). In the training process, Gumbel softmax is utilized to differentiate quantized features. Its calculation principle is as follows:

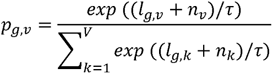

**Figure 3.**
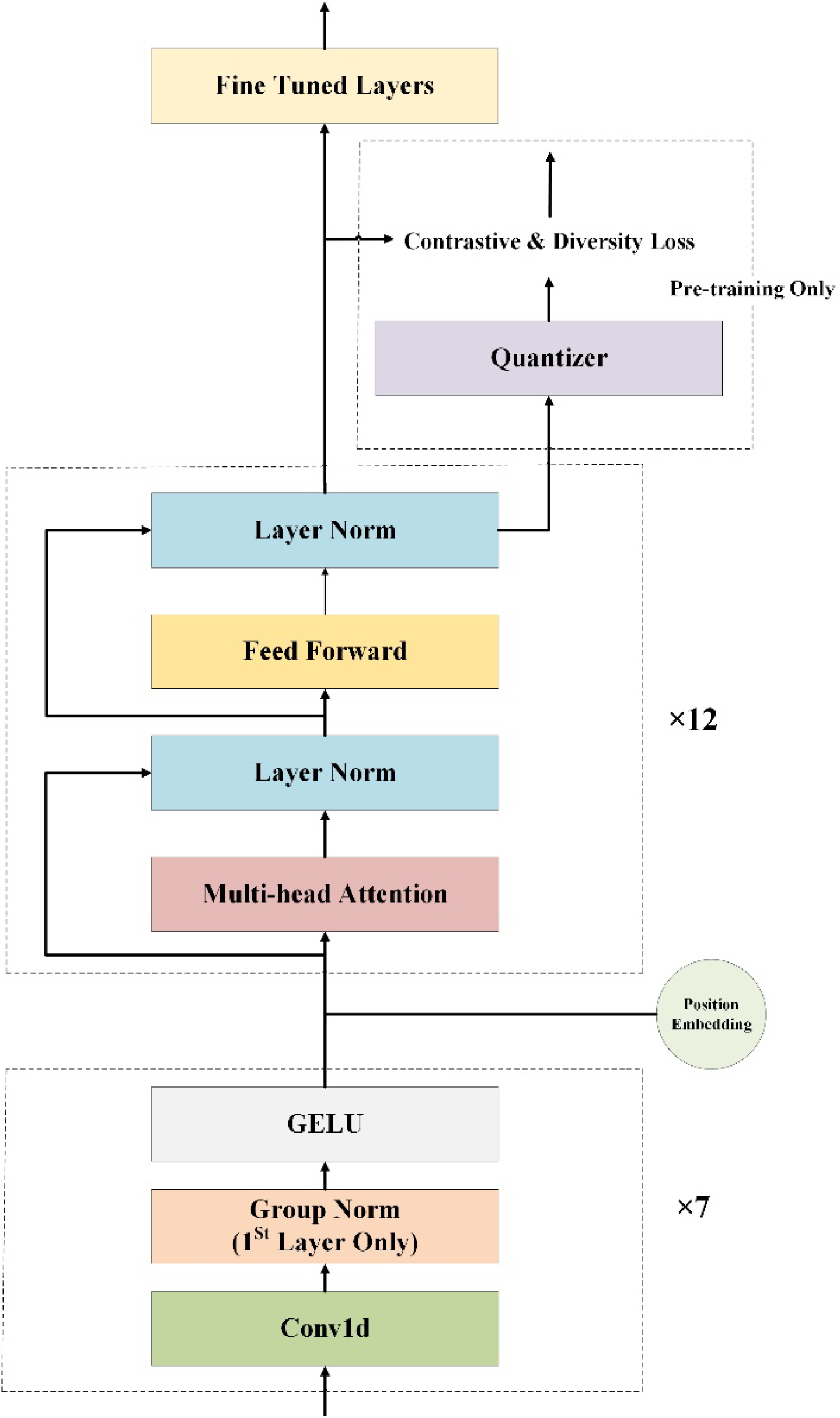
Self-supervised large-scale model architecture in BaseNet. The schematic illustrates the self-supervised large-scale model architecture developed in BaseNet. The model comprises three key components: a feature extraction module, an encoder module for context modeling, and a quantization module for learning discrete common features through self-supervised pre-training.

Where *τ* is a non-negative constant, *g*∈*G*, *v*∈*V*, *G* is the number of codebooks, and *V* is the dimension of each codebook.

The model achieves convergence through Contrastive Loss, which calculates the cosine similarity between context representation and the quantized representation, and Diversity Loss, which expands the space range of codebook. Their calculation principle are as follows:

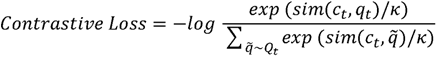

*c*_t_ represents the output feature of the encoder at the *t*-th masked time step, *q*_t_ denotes the discrete feature encoded by the quantization module at the *t*-th masked time step, and *Q*_t_ includes *q*_t_ and the *k* distractors at other time steps encoded by the quantization module.

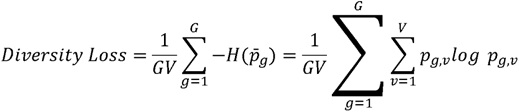

Through the aforementioned pre-training process, the encoder acquires generalized features with high robustness. In the fine-tuning phase, we devised seven strategies to enhance the model’s performance: 1). Adding a linear projection; 2). Incorporating an encoder layer followed by a linear projection; 3). Introducing two encoder layers followed by a linear projection; 4). Including three encoder layers followed by a linear projection; 5). Combining a linear layer, an encoder layer and a linear projection; 6). Including a linear layer, two encoder layers and a linear projection; and finally 7). Incorporating a linear layer, three encoder layers and a linear projection atop the models. These strategies are evaluated to determine the most suitable fine-tuning method for decoding nanopore sequencing signals. During fine-tuning, the models were optimized by minimizing the CTC loss.

### Joint loss training and rescore mechanism

The rescore model comprises three main components: a shared encoder, a CTC decoder and attention decoder, as shown in Figure 4. The shared encoder consists of 6 transformer encoder layers. The CTC decoder consists of a single linear layer, while the attention decoder consists of either three layers each of forward and reverse decoder layers (referred to as bidirectional decoder) or just three layers of forward decoder layers (referred to as unidirectional decoder) ^[25]^.

**Figure 4.**
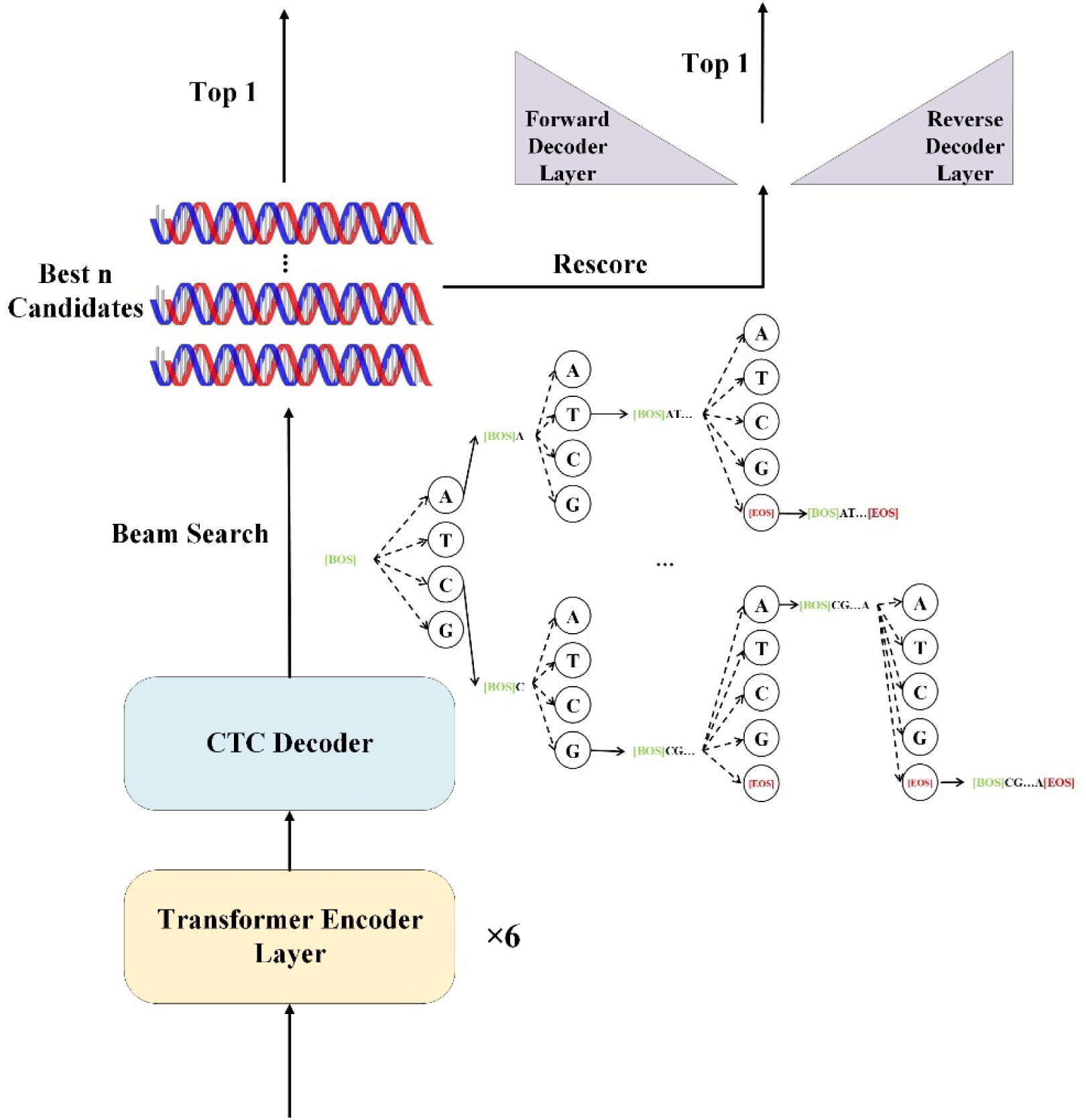
Rescore and joint loss training model in BaseNet. This schematic illustrates the rescore and joint loss training model used in BaseNet. The model consists of three main components: a shared encoder, a CTC decoder and attention decoders. The training process utilized a joint loss to optimize performance.

We train the model by converging a joint loss consisting of CTC loss and AED loss.

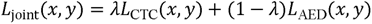

Where *x* is the output probability matrix, *y* is the label, and λ is a hyperparameter between 0 and 1.

During the rescore decoding stage, the CTC decoder first utilizes CTC prefix beam search to generate *n*-best candidate sequences. Subsequently, the attention decoder rescores the candidate sequences and selects the sequence with the highest score as the final decoding output.

### Paraformer

The Paraformer model includes an encoder for generating hidden representations, a predictor that generates acoustic embeddings and predicts sequence lengths, a sampler that randomly samples acoustic and target embeddings to create semantic embeddings, and a decoder that generates outputs based on the semantic embedding and hidden representations ^[26]^ (Figure 5).

**Figure 5.**
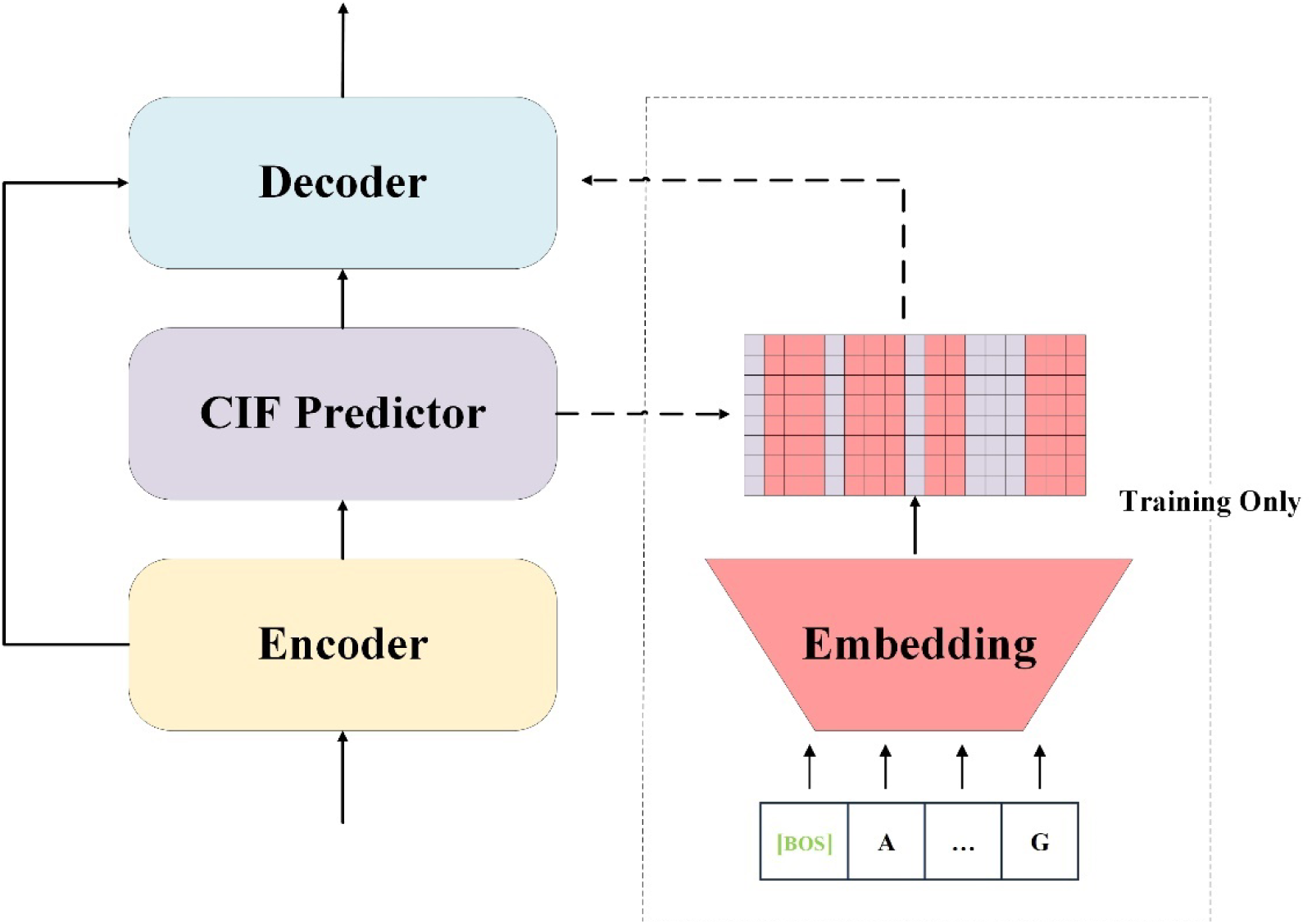
Paraformer Architecture in BaseNet. The schematic depicts the architecture of the Paraformer model developed in BaseNet. The model features an encoder for generating hidden representations, a predictor for producing acoustic embeddings and predicting sequence lengths, a sampler for randomly creating semantic embeddings, and a decoder for generating outputs.

However, due to extensive computational requirements, each epoch of training takes over 16 hours, we did not train this model because of the limitation on computational resources. Instead, the algorithm code is provided for reference and further exploration.

### Model Training

The AdamW optimizer was used to train the models, which dynamically adjusts the learning rate to optimize the training process. BaseNet provides three learning rate schedulers: CosineDecay^[31]^, Noam^[15]^ and WarmupLR^[32]^. Figure 6a illustrates the learning rate variation under different schedulers. Their calculation methods are as follows:

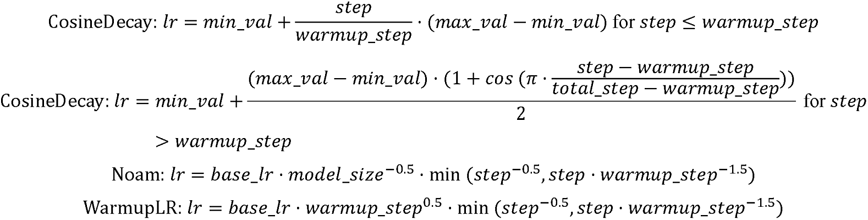

**Figure 6.**
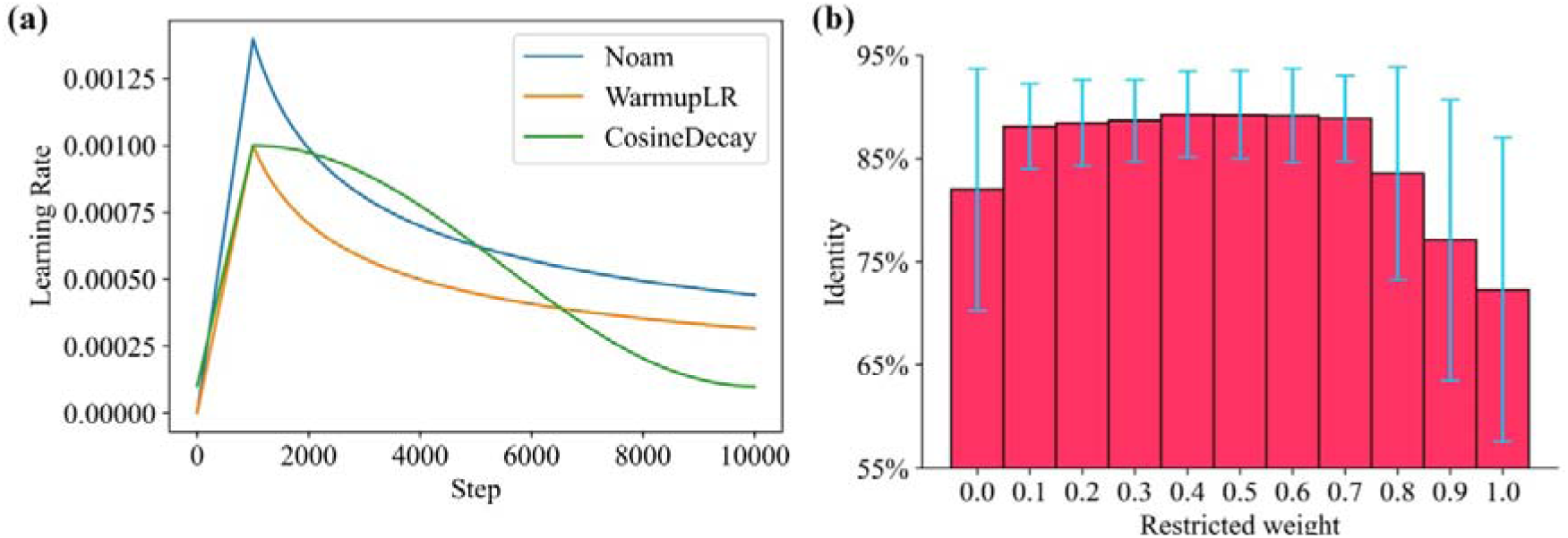
**Performance of BaseNet Under different learning rates and restricted weights.** (a) Learning rate variation under three schedulers: CosineDecay, Noam, and WarmupLR. The Warmup step is set to 1000 and the total step to 10000. (b) Autoregressive transformer prediction performance with different restricted weights. Performance significantly reduces with no EOS constraint (*w*=0) or strict constraints (*w*>=0.8). Optimal performance is observed within the range of *w* values from 0.1 to 0.7.

The training was performed on 8 Nvidia A100 40G GPUs. The autoregressive transformer (39,552,533 parameters) was trained using the cross-entropy loss function and Noam scheduler, excluding the loss for the padding token (PAD). The label sequences are smoothed with a smoothing coefficient of 0.1. The model was trained for 16 epochs with a batch size of 2. The training was performed in parallel, taking a total of 164 hours. Each of the large-scale fine-tuned models (97,137,664 parameters) was trained for 10 epochs with a batch size of 5, taking 124 hours through CosineDecay scheduler. The joint loss training by WarmupLR scheduler with bidirectional decoder (29,700,511 parameters) was performed 50 epochs, consuming 240 hours totally. The joint loss training by WarmupLR scheduler with unidirectional decoder (21,414,938 parameters) was performed 35 epochs, consuming 130 hours totally.

### Accuracy Evaluation

The basecalling results are aligned to the reference genome using minimap2^[33]^ and the prediction accuracy is calculated as the similarity between the basecalled sequence and the corresponding true sequence. The similarity is defined based on the following four metrics:

Identity: the percentage of matching bases between the basecalled sequence and the true sequence. It represents the proportion of correctly identified bases.

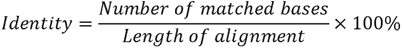

Mismatch rate: the percentage of bases in the basecalled sequence that do not match the true sequence. It represents the rate of incorrectly identified bases.

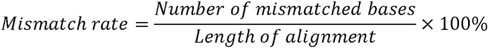

Insertion rate: the percentage of bases present in the basecalled sequence but absent in the true sequence. It represents the rate of false-positive insertions.

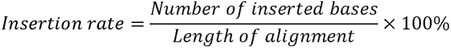

Deletion rate: the percentage of bases present in the true sequence but not detected in the basecalled sequence. It represents the rate of false-negative deletions.

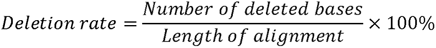

The overall median values of the above metrics were used to compare our model with other methods, which were also adopted in multiple basecaller studies for performance evaluation and comparation.

## Results and Discussion

### Autoregressive Termination Constraint

In autoregressive decoding, it is crucial to appropriately constrain the generation of the end of sequence (EOS) token to achieve optimal model performance. Here, we introduce a parameter called the constraint weight (*w*). The value of *w* ranges from 0 to 1. *w*=0 indicates no constraint on the generation of the EOS token, and *w*=1 means that the model does not consider generating the EOS token and decode until it reaches the maximum length. We compared the prediction accuracy of autoregressive transformer under different values of *w* on the same test data (Figure 6b). The results indicate that the model performance significantly deteriorates when there is no constraint on EOS generation (*w*=0) or when there is a strict constraint (*w*>=0.8). The model performs stably and well within the range of 0.1 to 0.7.

### Mapping between current signal and base sequence in the cross-attention layer

The cross-attention mechanism enables the decoder to derive hidden representations of the electrical signals from the encoder’s output and apply them to the base sequence decoding process. In the cross-attention layer, the queries *Q* come from the base sequences, which serve as the input to the decoder, while the keys *K* and values *V* are derived from the encoder’s output, originating from the current signals. This mechanism establishes attention relationships between the generated base sequences and the current signals.

At each position in Q, the cross-attention mechanism calculates the attention weights between the current base and all positions in the encoder’s output. This allows the decoder to focus on and utilize the corresponding current waveform information related to the base sequence. In the temporal domain, there is theoretically a linear correspondence between the current signals and the base sequences. Specifically, changes in the base sequences result in corresponding changes in current signals, and these changes are approximately linear.

Next we investigate whether transformer models can learn and capture this correlated relationship. To achieve this, attention weights were extracted and visualized from the cross-attention layers in autoregressive transformer (Figure 7). It is revealed that the strength of the linear relationship varies across different layers of the transformer decoder. Specifically, the relationship exhibits a weak-strong-weak pattern, with the highest strength observed in the fourth to sixth layers, indicating these layers are most effective at capturing the linear correlation. Conversely, the linear relationship is weaker or absent in the first to third layers and the seventh to eighth layers.

**Figure 7.**
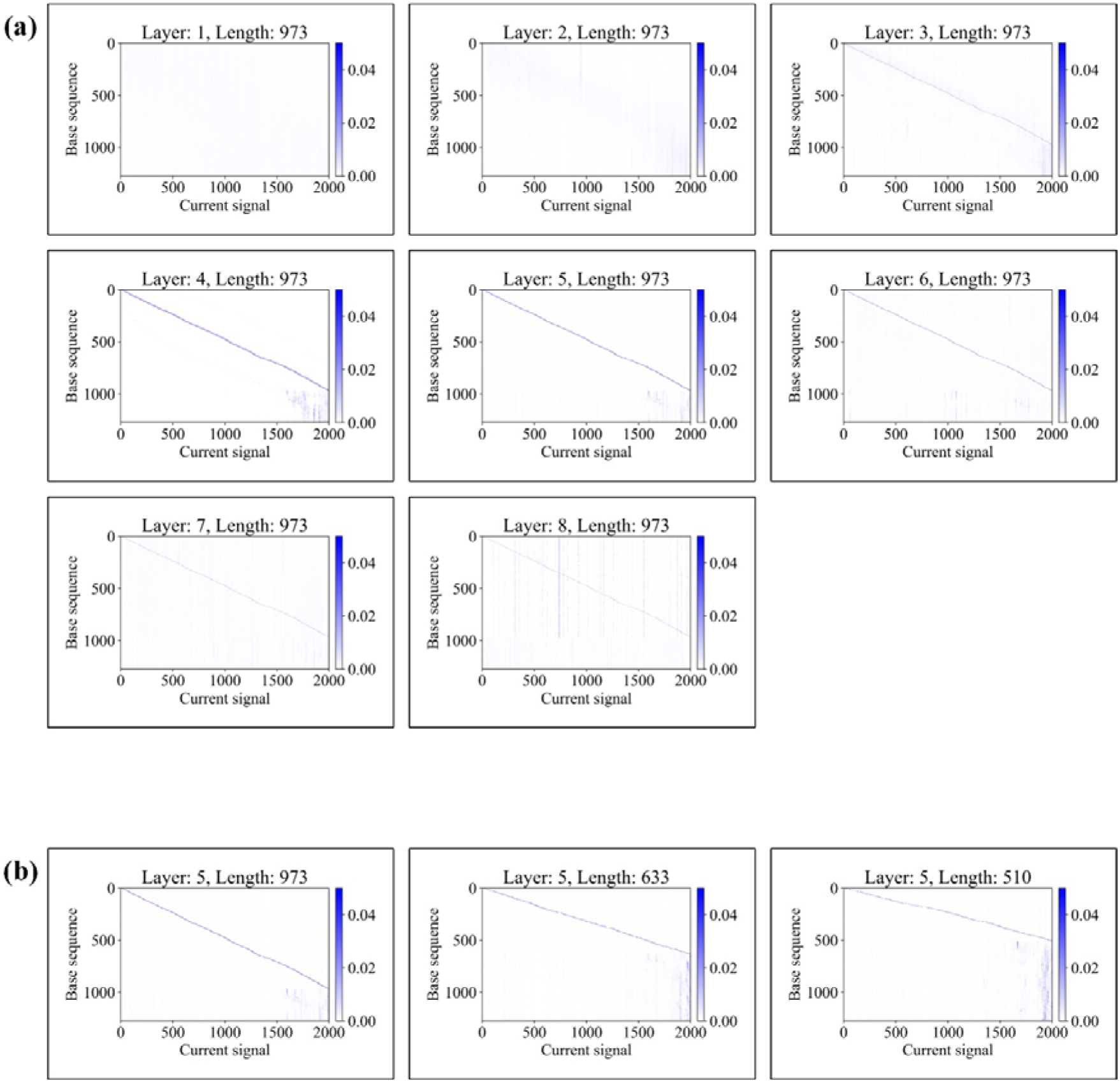
**Cross-attention weights between current signals and base sequences in transformer decoder.** (a) Visualization of cross-attention weights across different decoder layers for a specific sequence. The linear relationship between signal and sequence follows a weak-strong-weak pattern among layers. Layers four to six exhibit the strongest linear relationships, indicating that they effectively capture both local and global features. In contrast, layers one to three and seven to eight show no or weaker linear relationships, indicating a focus on either local or global features without integrating both. (b) Visualization of cross-attention weights across different sequences in decoder layer five.

The lack or weakening of the linear relationship in the first to third layers can be attributed to their focus on local features and limited ability to capture the overall linear relationship. These layers may primarily concentrate on extracting local patterns and features, resulting in a weaker capacity to learn the global linear relationship. The weakening of the linear relationship in the seventh and eighth layers can be attributed to their emphasis on global features while overlooking the local linear relationship. Both layers may excel in capturing global patterns and features but exhibit a weaker ability to learn the local linear relationships.

In contrast, the fourth to sixth layers demonstrate the strongest linear relationship, suggesting that they strike a better balance in attention weight learning, allowing them to simultaneously capture both local and global features. These layers appear to be more proficient in capturing the linear relationship between the base sequences and the current signals.

Thus, it is suggested that the attention weights from the fourth to sixth of the cross-attention layers can be extracted to establish an appropriate weight threshold. By doing so, the current signal points that exhibit a strong correlation with specific bass can be identified. This approach enables the assignment of current signal points to their corresponding bases, thereby establishing the correspondence between current signal points and bases to achieve alignment. This alignment method, based on attention weights and thresholds, offers a new strategy for data chunking and training set construction.

### Performance of large-scale model pre-training and fine-tuning

In this study, the large-scale model underwent five rounds of pre-training based on nanopore sequencing signals, followed by fine-tuning (signal fine-tuning). To determine the optimal fine-tuning strategy, we loaded the pre-training weights of wav2vec2.0 into our large-scale model and performed fine-tuning training (speech fine-tuning). After 10 epochs, we found that the model with only one added linear layer exhibited the best decoding performance, achieving 95.86% identity (Figure 8a). The finding indicates that incorporating too many or overly complex fine-tuning layers in large-scale pre-trained models does not necessarily enhance performance. Instead, adding just a linear layer based on the requirements of the task can yield desirable results. Excessive or complex fine-tuning layers may introduce too many parameters and complexity, leading to overfitting or performance degradation. Therefore, in subsequent fine-tuning studies, we opted to add only one linear layer.

**Figure 8.**
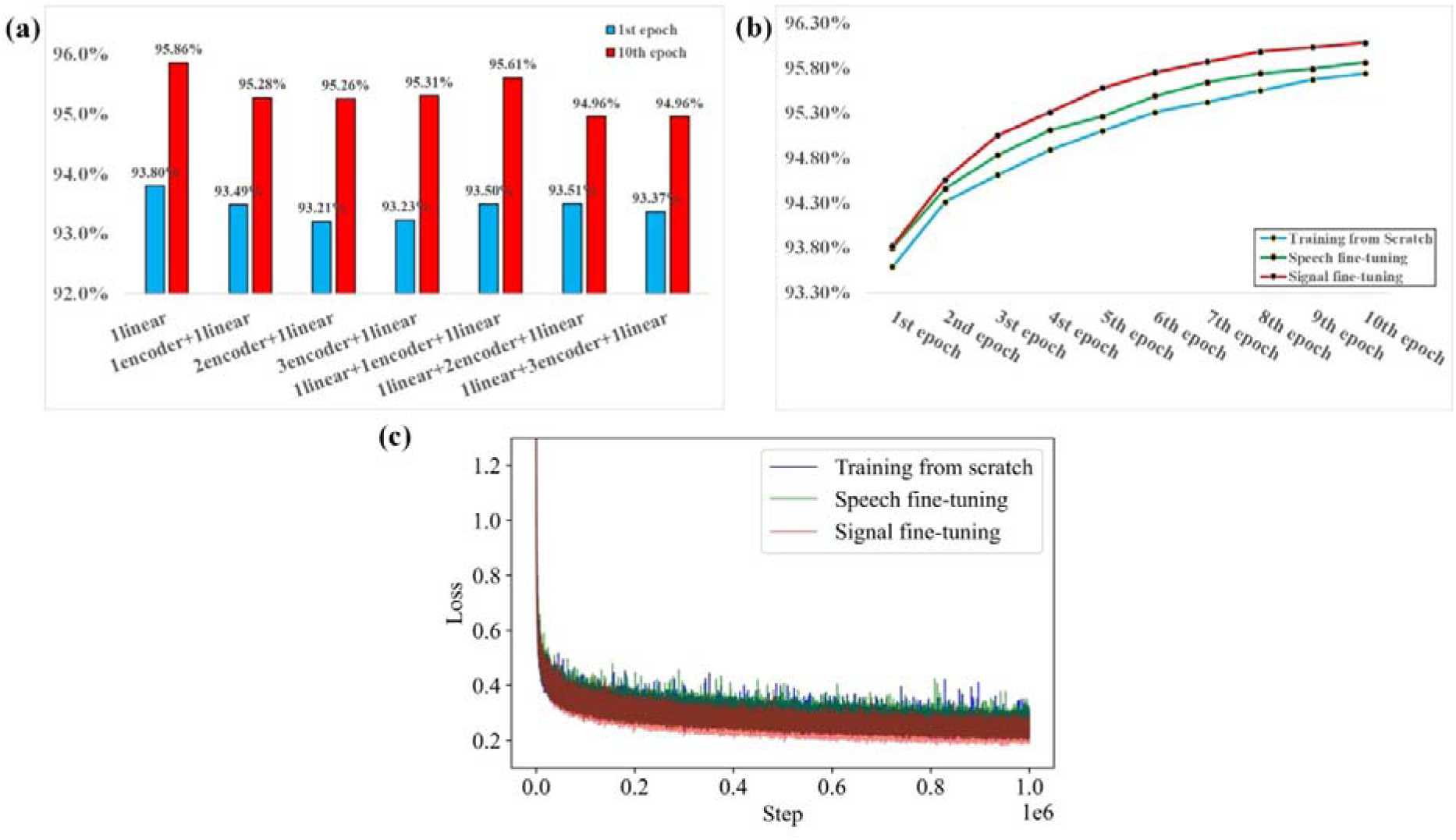
**Performance comparison of different large-scale models.** (a) Performance comparison of different fine-tuned models. Among the 7 fine-tuned models, the model with one additional linear layer achieved the best performance, reaching 95.86% identity after 10 epochs of fine-tuning. (b) Accuracy comparison of large models under different training conditions as indicated. (c) Training loss curves for signal fine-tuning, speech fine-tuning, and training from scratch.

Simultaneously, we also trained the large model from scratch using CTC loss (training from scratch). The results (Figure 8b and 8c) indicate that signal fine-tuning yields the best performance. This is understandable, as during the pre-training process, the large-scale model learns efficient, robust, and generic high-dimensional features from the input signals. Fine-tuning with supervised data refines these ‘generic features’ into ‘task-specific features’, thereby effectively applying these features to downstream tasks. Notably, the evaluation accuracy of speech fine-tuning significantly surpasses that of training from scratch, despite having similar training curves.

In speech recognition, the model extracts speech features from the original audio signal, while in nanopore sequencing current decoding, it extracts base sequence information from the current signal. These two tasks share similarities in signal extraction processes. Additionally, both speech recognition and nanopore sequencing current decoding require the model to recognize and learn specific patterns in the data.

These findings suggest shared underlying structures or patterns between speech recognition and nanopore sequencing current decoding. The high-dimensional hidden representations for both tasks exhibit similarities within the model, indicating that current signals and speech waveforms share common ‘generic features’.

### Joint loss training improves model performance

Next, we compared the performance of unidirectional and bidirectional decoder rescore models in the task of nanopore sequencing signal decoding. The models were evaluated at the 35^th^ epoch, with the unidirectional decoder achieving an identity of 94.57%, while the bidirectional decoder slightly outperformed it at 94.66% (Supplemental Table 1). Both models were trained using a combination of weighted CTC and AED loss (joint loss), enabling independent decoding by the encoder.

To further investigate the decoding capabilities of the encoder, we conducted ablation experiments by comparing the performance of the encoder-only approach with the encoder-decoder architecture. Surprisingly, both the unidirectional and bidirectional decoders achieved comparable performance to their respective complete models when using the encoder-only approach (Supplemental Table 1).

Subsequently, we focused on training the encoder alone using the CTC loss. The results revealed that training the encoder solely with CTC loss resulted in 94.18% identity after 35^th^ epoch (Supplemental Table 1).

These findings suggest that the non-autoregressive transformer rescore mechanism has limited potential for performance improvement in nanopore sequencing signal decoding. However, joint loss training of the model using CTC and attention mechanisms proves to be highly effective in enhancing model performance.

### Comparison of Different Basecallers

We compared the performance of five different basecallers, which are deep learning-based decoding models for sequencing data, in terms of decoding accuracy and inference speed. These basecallers include the Fine-tuned model, which is a large-scale pre-trained model fine-tuned with one linear layer, the Joint-CTC model trained through joint loss and bidirectional decoder, the Bonito’s CRF model, the Fast-CRF model which replaces LSTM layers of Bonito-CRF model with fast attention layers, and the latest third party open source basecaller SACall.

The inference was conducted on an NVIDIA RTX 3090 24G GPU. As shown in Supplemental Tables 2 and 3, the Fine-tuned model outperformed the latest ONT basecaller in decoding accuracy after only 10 epochs of training. Although the performance of Joint-CTC model and the Fast-CRF model slightly lag behind the Bonito-CRF model, they significantly surpass SACall. These findings establish that BaseNet achieves reasonable performance and compares favorably with the ONT basecaller. Thus, BaseNet provides the state-of-the-art open source basecallers.

## Conclusion

We introduce BaseNet, an open-source toolkit for nanopore sequencing signal decoding based on state-of-the-art transformer algorithm. Experiments and comparisons between BaseNet and other basecallers demonstrate that BaseNet achieves reasonable performance and comparable results. In addition, our study reveals several insights: cross-attention weights of transformer effectively map the correspondence between current signals and base sequences; joint loss training with the addition of forward and reverse decoders aids in better model convergence; large-scale pre-trained model achieve higher decoding accuracy; and there are common ‘generic features’ between speech waveforms and sequencing signals in model representation.

## Acknowledgements

This work is partly supported by grants from the Ministry of Science and Technology of China (2019YFA0707001 and 2021YFF0700201) and the Strategic Priority Research Program of the Chinese Academy of Sciences (XDB37020102).

## Data and code availability

The data of this study was taken from the study of Wick et. al (https://doi.org/10.1099/mgen.0.000132.).

The code, model weights of the Pre-trained and Fine-tuned large-scale model, Joint-CTC model, and Fast-CRF model to BaseNet can be accessed from Github (https://github.com/liqingwen98/BaseNet).

