## Supplemental Table 1-3 for "BaseNet: A Transformer-Based Toolkit for Nanopore Sequencing Signal Decoding"

**Supplemental Table 1. Performance comparison of joint training models using different decoding methods.**

|  | Bi-decoder | Uni-decoder | w/o Decoder |
| --- | --- | --- | --- |
| Rescore | 94.66% | 94.57% | Nan |
| CTC | **94.70%** | 94.62% | 94.18% |

**Supplemental Table 2. Comparison of decoding performance of different basecallers.**

| Basecaller | Identity (%) | Deletion rate (%) | Insertion rate (%) | Mismatch rate (%) | Speed (E+6 / s) |
| --- | --- | --- | --- | --- | --- |
| **Fine-tuned model** | **94.37** | **1.36** | **1.09** | **3.14** | 0.43 |
| Bonito-CRF model | 93.35 | 1.64 | 1.10 | 3.88 | **9.48** |
| **Joint-CTC model** | 93.32 | 1.82 | 1.21 | 3.60 | 4.09 |
| **Fast-CRF model** | 91.76 | 2.00 | 1.57 | 4.65 | 8.07 |
| SACall | 90.45 | 2.42 | 2.07 | 4.98 | 0.97 |

**Supplemental Table 3. Decoding performance of different basecallers on different species.**

| Species | Basecaller | Identity (%) | Insertion rate (%) | Deletion rate (%) | Mismatch rate (%) |
| --- | --- | --- | --- | --- | --- |
| Acinetobacter Pittii | Fine-tuned model | **94.71** | 1.08 | **1.37** | **2.85** |
|  | Bonito_CRF | 94.64 | **0.89** | 1.44 | 3.04 |
|  | Joint-CTC model | 93.66 | 1.22 | 1.76 | 3.34 |
|  | Fast-CRF model | 92.04 | 1.57 | 2.03 | 4.38 |
|  | SACall | 90.50 | 2.29 | 2.29 | 4.91 |
| Stenotrophomonas Maltophilia | Fine-tuned model | **93.93** | **1.07** | **1.47** | **3.48** |
|  | Bonito_CRF | 92.29 | 1.21 | 1.93 | 4.51 |
|  | Joint-CTC model | 93.06 | 1.20 | 1.82 | 3.86 |
|  | Fast-CRF model | 91.08 | 1.58 | 2.21 | 5.08 |
|  | SACall | 89.98 | 2.60 | 2.00 | 5.30 |
| Shigella Sonnei | Fine-tuned model | **93.63** | 1.32 | **1.48** | **3.55** |
|  | Bonito_CRF | 93.22 | **1.21** | 1.56 | 3.99 |
|  | Joint-CTC model | 92.42 | 1.36 | 2.09 | 4.06 |
|  | Fast-CRF model | 91.36 | 1.75 | 1.94 | 4.93 |
|  | SACall | 90.28 | 1.46 | 3.02 | 5.14 |
| Haemophilus Haemolyticus | Fine-tuned model | **95.80** | **0.94** | **1.10** | **2.11** |
|  | Bonito_CRF | 94.95 | **0.94** | 1.24 | 2.82 |
|  | Joint-CTC model | 94.56 | 1.10 | 1.61 | 2.69 |
|  | Fast-CRF model | 93.03 | 1.35 | 1.93 | 3.66 |
|  | SACall | 89.38 | 3.60 | 1.61 | 5.29 |
| Klebsiella Pneumoniae Nuh29 | Fine-tuned model | **94.59** | **0.97** | **1.34** | **3.04** |
|  | Bonito_CRF | 92.56 | 1.19 | 1.79 | 4.37 |
|  | Joint-CTC model | 93.66 | 1.12 | 1.71 | 3.44 |
|  | Fast-CRF model | 91.86 | 1.43 | 2.08 | 4.58 |
|  | SACall | 90.35 | 2.81 | 1.81 | 4.91 |
| Klebsiella Pneumoniae Ksb2 | Fine-tuned model | **93.90** | **1.16** | **1.48** | **3.43** |
|  | Bonito_CRF | 92.08 | 1.26 | 1.95 | 4.66 |
|  | Joint-CTC model | 92.82 | 1.27 | 1.96 | 3.90 |
|  | Fast-CRF model | 91.30 | 1.60 | 2.12 | 4.95 |
|  | SACall | 90.09 | 2.24 | 2.55 | 5.18 |
| Klebsiella Pneumoniae Inf042 | Fine-tuned model | **94.09** | **1.13** | **1.43** | **3.31** |
|  | Bonito_CRF | 92.30 | 1.22 | 2.02 | 4.47 |
|  | Joint-CTC model | 93.01 | 1.25 | 1.90 | 3.78 |
|  | Fast-CRF model | 91.36 | 1.56 | 2.24 | 4.83 |
|  | SACall | 90.37 | 2.26 | 2.49 | 5.01 |
| Klebsiella Pneumoniae Inf032 | Fine-tuned model | **93.89** | **1.14** | **1.48** | **3.45** |
|  | Bonito_CRF | 92.41 | 1.25 | 1.87 | 4.42 |
|  | Joint-CTC model | 92.77 | 1.25 | 1.99 | 3.91 |
|  | Fast-CRF model | 91.50 | 1.58 | 2.08 | 4.81 |
|  | SACall | 90.61 | 1.47 | 2.86 | 4.96 |
| Serratia Marcescens | Fine-tuned model | **94.52** | 0.97 | **1.30** | **3.17** |
|  | Bonito_CRF | 93.86 | **0.89** | 1.51 | 3.69 |
|  | Joint-CTC model | 93.57 | 1.11 | 1.68 | 3.60 |
|  | Fast-CRF model | 91.53 | 1.57 | 1.91 | 4.96 |
|  | SACall | 90.72 | 2.28 | 2.01 | 4.94 |
| Staphylococcus Aureus | Fine-tuned model | 95.71 | 0.96 | 1.12 | 2.21 |
|  | Bonito_CRF | **96.40** | **0.68** | **0.96** | **1.95** |
|  | Joint-CTC model | 94.75 | 1.06 | 1.54 | 2.62 |
|  | Fast-CRF model | 93.66 | 1.39 | 1.66 | 3.31 |
|  | SACall | 91.64 | 1.70 | 2.47 | 4.18 |
